# Distinct developmental phenotypes result from mutation of Set8/KMT5A and histone H4 lysine 20 in *Drosophila melanogaster*

**DOI:** 10.1101/2022.01.30.478405

**Authors:** Aaron T. Crain, Stephen Klusza, Robin L. Armstrong, Priscila Santa Rosa, Brenda R. S. Temple, Brian D. Strahl, Daniel J. McKay, A. Gregory Matera, Robert J. Duronio

## Abstract

Mono-methylation of histone H4 lysine 20 (H4K20me1) is catalyzed by Set8/KMT5A and regulates numerous aspects of genome organization and function. Loss-of-function mutations in *Drosophila melanogaster Set8* or mammalian *KMT5A* prevent H4K20me1 and disrupt development. Set8/KMT5A also has non-histone substrates, making it difficult to determine which developmental functions of Set8/KMT5A are attributable to H4K20me1 and which to other substrates or to non-catalytic roles. Here, we show that human KMT5A can functionally substitute for Set8 during *Drosophila* development and that the catalytic SET domains of the two enzymes are fully interchangeable. We also uncovered a role in eye development for the N-terminal domain of Set8 that cannot be complemented by human KMT5A. Whereas *Set8*^*null*^ mutants are inviable, we found that an R634G mutation in the SET domain predicted to ablate catalytic activity resulted in viable adults, suggesting important non-catalytic functions of Set8. Similarly, flies that were engineered to express only unmodifiable H4 histones (*H4*^*K20A*^) can also complete development, but they are phenotypically distinct from *H4*^*K20R*^, *Set8*^*null*^, and *Set8*^*R634G*^ animals. Taken together, our results demonstrate functional conservation of KMT5A and Set8 enzymes, as well as distinct roles for Set8 and H4K20me1 in *Drosophila* development.

## Introduction

The formation of chromatin from DNA and histones regulates genome function and is critical for development of multicellular organisms. The post-translational modification (PTM) of histone N-terminal tails modulates the organization of chromatin and thereby helps regulate replication, repair, and transcription of the genome.^69^ Consequently, dysregulation of histone PTMs is thought to disrupt animal development. However, our understanding of how particular histone PTMs influence specific developmental processes is incomplete. For instance, methylation of histone H4 lysine 20 (H4K20me) has been implicated in the control of transcription,^2,9,14,17,38–40,43,44,46,78,90,97,99,105^ DNA replication and repair,^9,12,14,26,32,34,42,79,100^ chromosome condensation during mitosis,^9,14,16,40^ and heterochromatin assembly,^9,14,61,79,84^ but the requirement for these putative H4K20me functions has not been directly interrogated during animal development.^49^

In most animal genomes, H4K20 mono-methylation (H4K20me1) is catalyzed by a conserved enzyme variably termed KMT5A/Set8/SETD8/PR-Set7 that contains a catalytic SET domain.^27,61^ Subsequent di- and tri-methylation of H4K20 is carried out by SET domain-containing Suv4-20 enzymes, of which there are two in mammals and one in *Drosophila*.^9,71,74,92,98^ Developmental roles for H4K20me are typically investigated by mutations that eliminate or alter the activity of these enzymes. Although most of this work has been done using knockdown methods in cell culture,^2,8,16,17,33,34,36,37,62,64,67,70,80,81,85,86,89,97^ a small number of studies were conducted using mutant animals.^7,27,40,44,63,72,75^ For instance, loss of the H4K20me2 methyltransferase Suv4-20h1 in mice causes early developmental defects, resulting in either embryonic or perinatal lethality.^75^ In contrast, animals that lack the H4K20me3 methyltransferase Suv4-20h2 develop normally.^75^ *Drosophila Suv4-20* null mutations display no overt developmental defects, suggesting no essential requirement for H4K20me2 and H4K20me3 in flies.^71^ In contrast, loss of H4K20 mono-methyltransferases causes severe developmental phenotypes: Fly *Set8 (*FlyBase annotation *PR-Set7; CG3307)* and mouse *KMT5A* null mutants are inviable and exhibit a developmental arrest that is accompanied by reduction of H4K20me and a variety of defects including smaller larval tissues in flies and increased apoptosis in mouse embryos.^34,40,63,72^ Mutant cells also have defects in cell cycle progression and accumulate DNA damage.^9,14,95^ These cellular and developmental defects have been attributed to loss of downstream functions that require H4K20 methylation. Consistent with this interpretation, a KMT5A R265G mutation predicted to abolish catalytic activity does not support embryonic development,^63^ suggesting that KMT5A catalytic activity is required for proper mouse development.

Each of these analyses is confounded by observations that Set8-family enzymes have protein substrates in addition to H4K20.^23,34,76,83^ Moreover, many of these other substrates, such as p53 and PCNA (Proliferating Cell Nuclear Antigen), regulate critical aspects of genome function.^23,76,83,93^ Finally, recent work from our group using engineered *Drosophila* histone mutant genotypes demonstrated that H4K20 is dispensable for DNA replication and organismal viability.^49^ Thus, the contributions of H4K20me to animal development are not fully determined.

Here, we compare phenotypes caused by mutation of *Set8* and *H4K20* in *Drosophila*. The data show that the essential function played by Set8 in fly development is either non-catalytic or is largely independent of its histone H4K20 methylation activity. We also demonstrate that human KMT5A can functionally substitute for loss of Set8 in *Drosophila*, indicating that flies can provide critical information about evolutionarily conserved functions of H4K20 mono-methyltransferases during development.

## Results

### Set8 is the appropriate designation for the *Drosophila* H4K20 mono-methyltransferase

The *Drosophila melanogaster* genome encodes fourteen SET [Su(var)3-9, Enhancer-of-zeste, Trithorax] domain lysine methyltransferases (FlyBase)^24,35,56,57,74,77^ (Figure 1). A related family of proteins, called PRDMs, is characterized by the presence of a PR domain [PRDF1 (positive regulatory domain I-binding factor 1) and RIZ1 (retinoblastoma protein-interacting zinc finger gene 1)] along with a variable number of Cys2-His2 (C2H2) zinc fingers.^88^ The PR domain is an evolutionarily recent subtype of the SET domain, although not all PRDMs encode active methyltransferases.^88^ There are four PRDM proteins (Blimp-1, Hamlet, CG43347, Prdm13) in insect genomes whereas this family has expanded to nineteen proteins in humans.^29,48,88^

**Figure 1.**
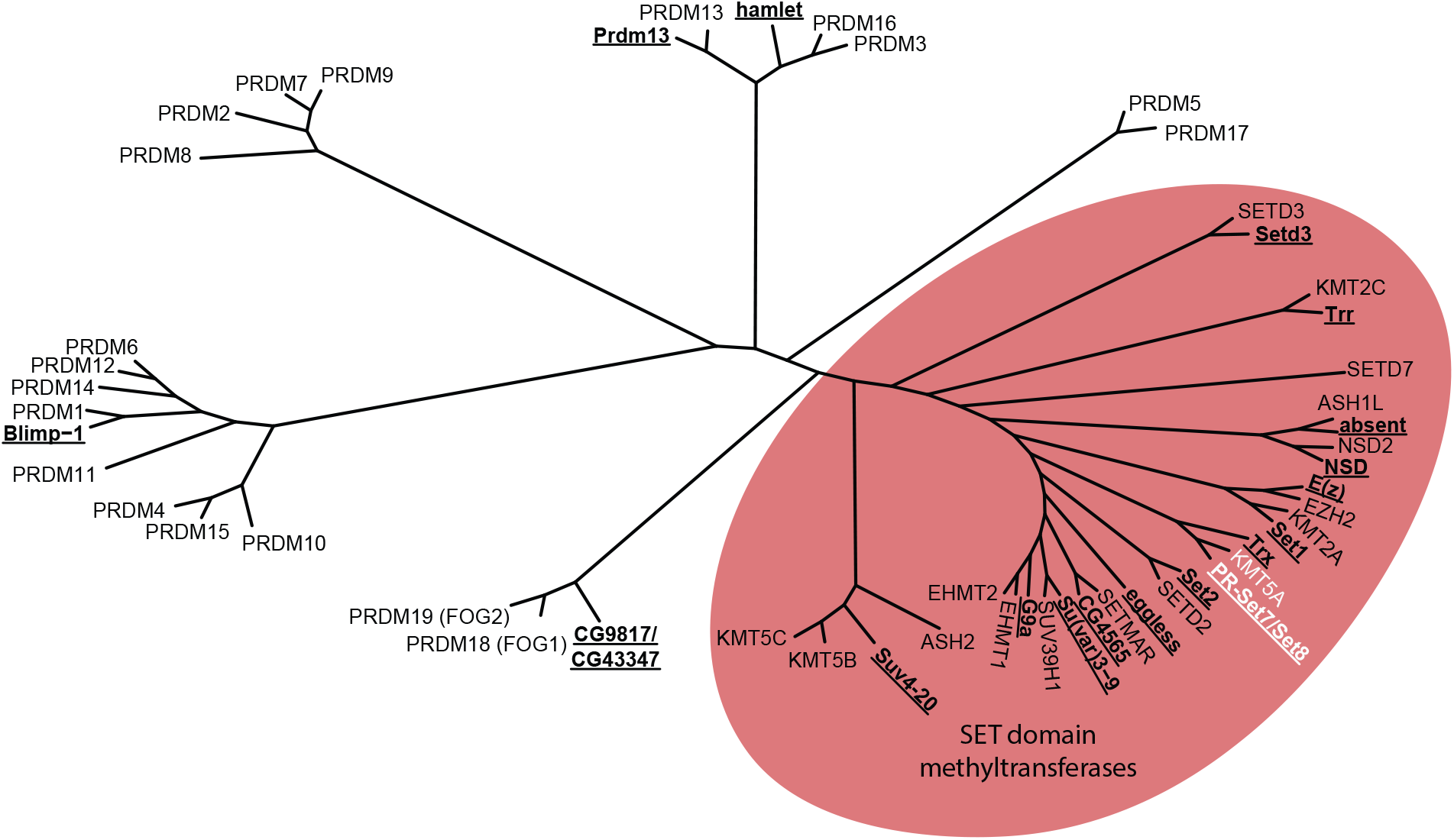
Evolutionary relationship of *Drosophila* and human SET and PRDM containing proteins. Unrooted tree produced from an alignment of human and *Drosophila* PRDM and SET domain family proteins using ClustalOmega. *Drosophila* proteins are indicated with bold, underlined text. The red oval highlights the grouping of SET domain family proteins, including PR-Set7/Set8 and KMT5A (white).

In *Drosophila*, the protein encoded by *PR-Set7/CG3307* is orthologous to the human H4K20 methyltransferase SETD8/KMT5A and it neither contains a PR domain nor any predicted zinc finger motifs^15,27,61^ (Figure 1). In contrast, human PRDM7 (PR/SET Domain 7) is an H3K4 methyltransferase that is most closely related to the KRAB and Zn finger domain protein, PRDM9^10^ (Figure 1). Moreover, human SETD7/Set7/Set9 is yet another human H3K4 methyltransferase^91^ distinct from *Drosophila* PR-Set7/CG3307 (Figure 1). To avoid further confusion, we propose to officially rename *CG3307* as *Set8* and refer to this protein as Set8 throughout the manuscript.

### Human KMT5A rescues loss of Set8 in *Drosophila*

KMT5A and Set8 are essential for the development of mice and flies, respectively, and mutating these enzymes results in defects in cell cycle progression, DNA damage response, and chromatin compaction in both organisms.^40,63,70^ The SET domains of Set8 and human KMT5A are 57% identical (Supplemental figure 1). Therefore, we hypothesized that KMT5A and Set8 perform the same biological functions in *Drosophila* and mammals and that human KMT5A would rescue loss of Set8 in *Drosophila*. To test this hypothesis, we engineered a *KMT5A* open reading frame that was codon-optimized for translation in *Drosophila* and expressed in the context of the native *Set8* gene (a 4774 bp genomic fragment including 1325 bp upstream of the ORF and 2021 bp downstream of the ORF including both native 5’ and 3’ UTRs). Using this engineered *KMT5A* allele, we generated transgenes located on the same chromosome as the *Set8*^*20*^ null allele^40^ (Figure 2A). Whereas *Set8*^*20/20*^ mutants die as early pupae, animals expressing KMT5A in a *Set8*^*20/20*^ background pupate normally and complete development at similar frequencies as wild type animals or *Set8*^*20/20*^ animals rescued with a control *Set8* transgene (Figure 2B, C). Although *Set8*^*20/20*^ animals rescued by KMT5A are viable and fertile, we observed a rough eye phenotype in 58% of adult flies (Figure 2E). The *Drosophila* compound eye is a highly organized tissue containing ∼800 photoreception structures termed ommatidia, each composed of eight photoreceptor neurons and a set of accessory cells. Many processes contribute to proper formation of the adult eye, including cell cycle progression, cell death, and ultimately cell differentiation. Disruption of any one of these processes can contribute to ommatidial irregularities that manifest as a visible “roughness” of the adult eye.^6,94^ Even subtle defects in gene functions required for eye development can result in rough eyes, and thus we conclude that KMT5A fully rescues most, but not all, Set8 functions in *Drosophila*.

**Figure 2.**
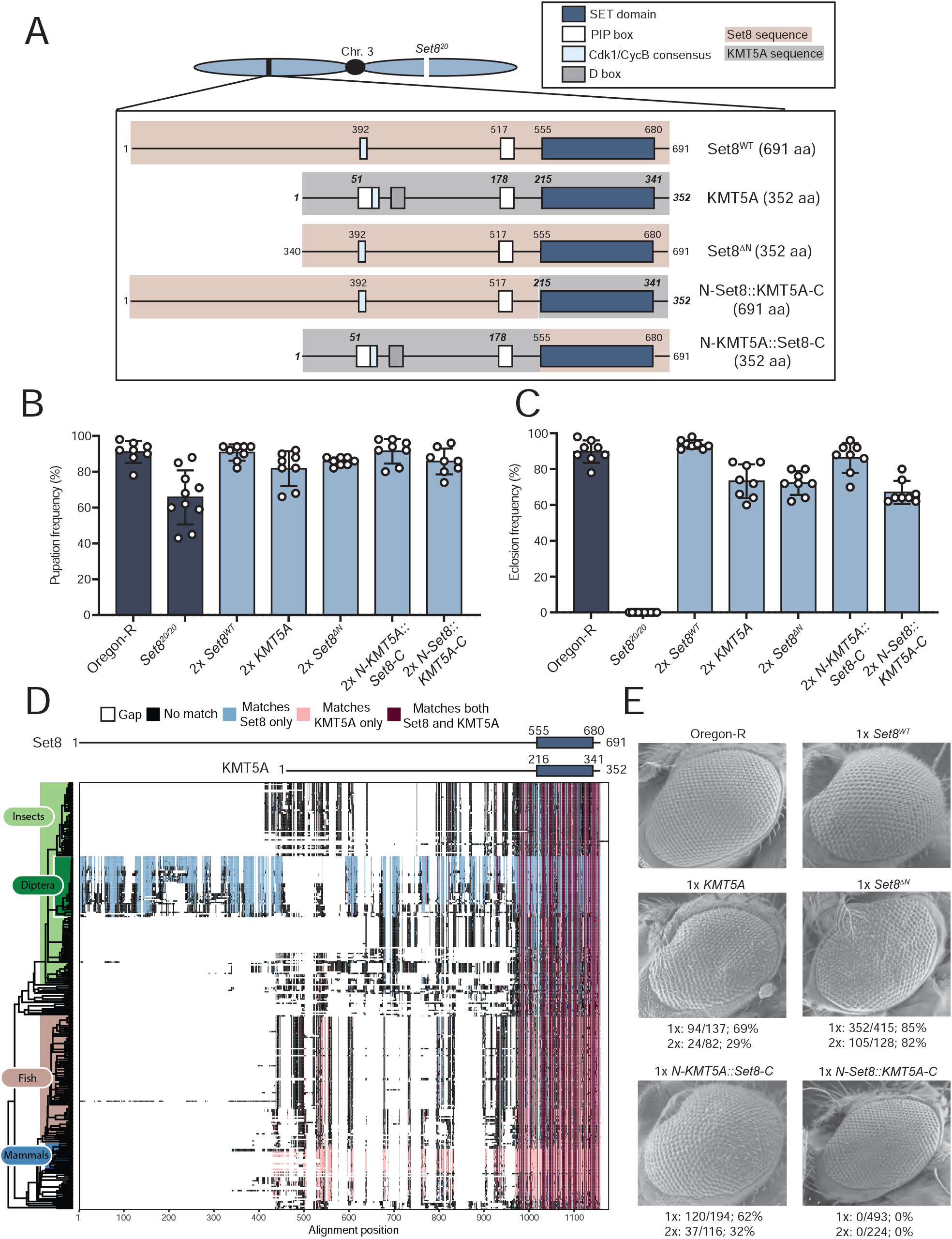
Human KMT5A functionally substitutes for Set8 during *Drosophila* development. A) Diagram of Set8, KMT5A, and Set8/KMT5A chimeric proteins expressed from transgenes located on chromosome 3, which also contains the *Set8*^*20*^ null allele. Red shading and non-bold, non-italic numbers indicate Set8 sequence. Gray shading and bold, italic numbers indicate KMT5A sequence. In parentheses is the total number of amino acids in each protein product. B) Pupation of *Set8*^*WT*^, *KMT5A*, and chimera genotypes. Each circle represents the percentage of 40-50 larvae in a vial that reached pupation. The mean and standard deviation of these percentages for 8-10 vials are shown for the indicated genotypes. All transgenic genotypes are in the *Set8*^*20/20*^ homozygous null background. “2x” indicates that each transgene is also homozygous. C) Eclosion into adults of *Set8*^*WT*^, *KMT5A*, and chimera genotypes. Here, each circle represents a vial of 40-50 larvae, and 8 vials for each of the indicated genotypes were scored. Genotypes are as in panel B. D) Annotated alignment of Set8-related proteins. 301 homologous proteins with over 50% identity to Set8 as identified via BLAST were aligned using Clustal Omega and ordered by phylogeny. Set8 and KMT5A schematics are shown at the top of the diagram with the SET domains indicated by dark blue boxes. Residues of each protein in the alignment that match both Set8 and KMT5A exactly are colored dark red. Those that match only Set8 are colored light blue, and those that match only KMT5A are colored pink. Residues that match neither are colored black. Gaps in the alignment are indicated by white space. E) SEM images of adult eyes of flies of the indicated genotypes. The penetrance of flies displaying a phenotype like that shown is indicated below each image. “1x” and “2x” indicate flies containing either 1 or 2 copies, respectively, of the transgene expressing Set8, KMT5A, or Set8/KMT5A chimeras in the *Set8*^*20/20*^ homozygous null background.

### The N-terminus of Set8 is dispensable for *Drosophila* viability but plays a role in eye development

Although the SET domains of Set8 and KMT5A are 57% identical, the full-length proteins are only 21% identical (Supplemental figure 1). The N-terminal region (554aa) of Set8 is predicted to be largely unstructured and is not well-conserved with KMT5A (Figure 2A). A multiple protein alignment of 301 BLAST hits with greater than 50% identity to the full-length Set8 protein revealed that this N-terminal region of Set8 is unique to flies (order Diptera), whereas the SET domain is highly conserved across all represented organisms (Figure 2D). To test whether the N-terminal region of Set8 functions in *Drosophila* development we engineered a transgene encoding a Set8 protein lacking the first 339 amino acids (*Set8*^*ΔN*^), which would produce a protein the size of KMT5A (Figure 2A). The *Set8*^*ΔN*^ transgene rescued *Set8*^*20/20*^ lethality resulting in fully viable and fertile adults with a highly penetrant (82%) rough eye phenotype (Figure 2B, C, E). Although we were unable to assess the protein accumulation of Set8^ΔN^ because the epitope recognized by the Set8 antibody is within the N-terminal region, these results indicate that the N-terminal 339 amino acids are dispensable for normal development except in the eye. To test whether the eye function could be provided by KMT5A, we generated a chimeric transgene with the Set8 N-terminus (1-554) fused to the KMT5A C-terminus (N-Set8::KMT5A-C) and a reciprocal chimeric transgene with the KMT5A N-terminus (1-214) fused to the Set8 C-terminus (N-KMT5A::Set8-C). Both transgenes fully rescued viability and fertility of *Set8*^*20/20*^ mutants (Figure 2 B, C). Further, *N-KMT5A::Set8-C* animals displayed a rough eye phenotype like *KMT5A* and *Set8*^*ΔN*^ animals (Figure 2E). By contrast, flies expressing the *N-Set8::KMT5A-C* chimera were fully viable and fertile with morphologically normal eyes, indicating the human KMT5A SET domain is functionally equivalent to that from *Drosophila* Set8 (Figure 2B, C). We conclude that the N-terminal 339 amino acids of Set8 are dispensable for *Drosophila* viability and fertility but have a function in eye development that cannot be provided by the first 214 amino acids of human KMT5A.

### A SET domain mutation predicted to block methyltransferase activity does not result in a Set8 null phenotype

Many of the established roles for the KMT5A/Set8 lysine methyltransferase have been attributed to its catalytic activity, primarily by using cell culture-based assays.^2,8,16,17,33,34,36,37,62,64,67,70,80,81,85,86,89,97^ To determine whether methyltransferase activity is required for Set8 function during *Drosophila* development, we engineered point mutations in the SET domain that are predicted to ablate catalytic activity^27,61^ (Figure 3A). SET domains are highly conserved and contain evolutionarily invariant residues within the catalytic core (Figure 3B). As shown in Figure 3C, two of these residues (R634 and H638) make critical contacts with the methyl donor, S-adenosyl methionine (SAM). Mutation of the homologous Arg residue in the human enzyme (R265) to Gly blocks methyltransferase activity *in vitro* using nucleosomal substrates, and this substitution has been used in numerous studies of Set8/KMT5A proteins to create catalytically inactive enzymes.^1,2,17,26,33,38,61,63,76,80,84,86,96^ We therefore engineered a *Set8*^*R634G*^ mutation (hereafter *Set8*^*RG*^) in the context of the rescuing genomic fragment used in the experiments above. We also engineered an R634G, H638L double mutation (hereafter *Set8*^*RGHL*^). Using *in silico* structural models based on the solved human KMT5A structure and molecular dynamics simulations (Figure 3C), each of these amino acid changes were evaluated for their impact on SAH binding and H4 peptide binding. While H4 peptide binding was minimally impacted, the mutations were shown to disrupt SAH binding and thus are predicted to reduce or eliminate methyltransferase activity of the mutant Set8 proteins (Figure 3C, D).

**Figure 3.**
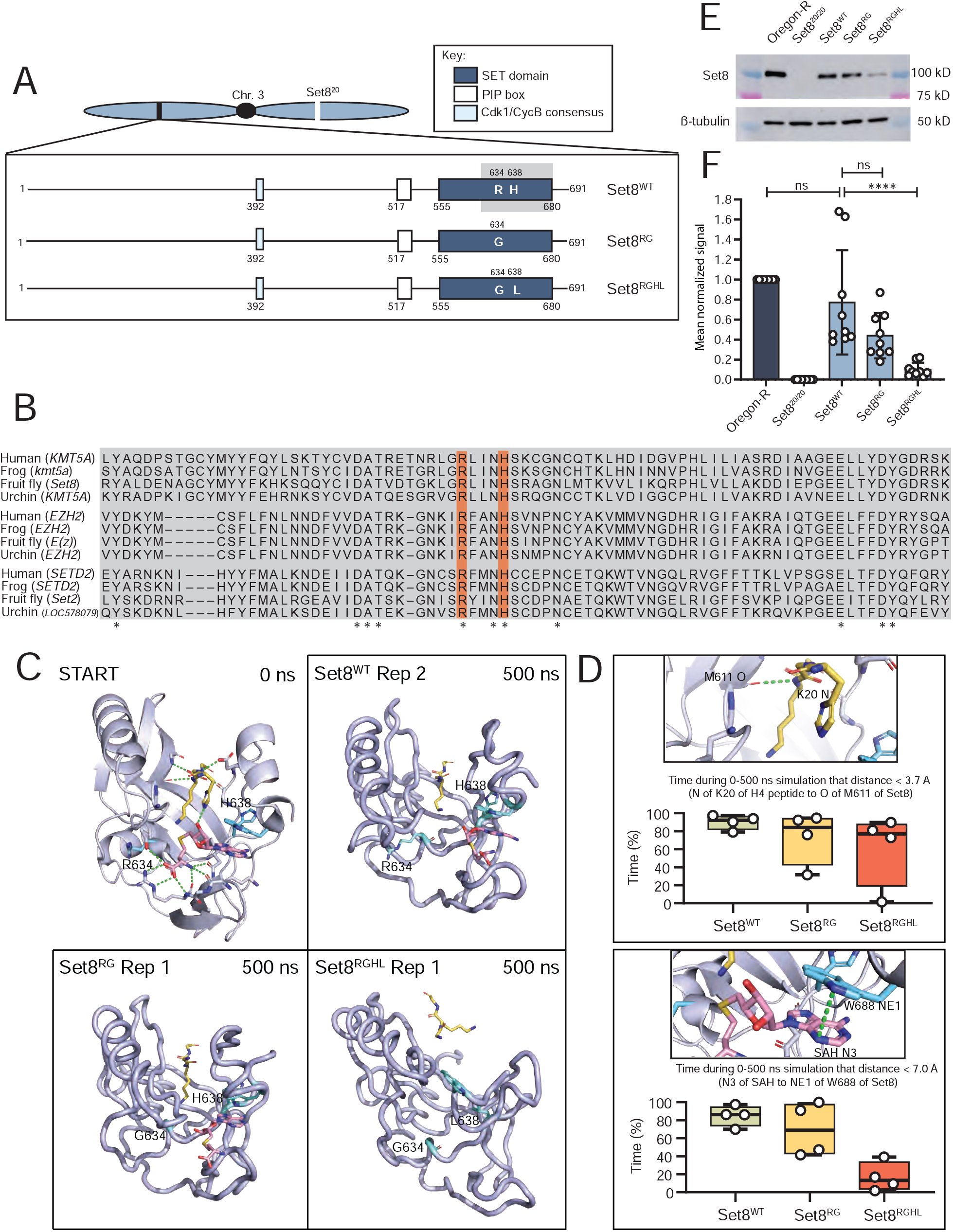
Generation Set8 proteins predicted to be catalytically inactive. A) Diagram of Set8(WT), Set8(RG) and Set8(RGHL) proteins expressed from transgenes located on chromosome 3. B) Conservation of the Set8 Arg634 and Leu638 residues (orange bars) among KMT5A proteins from human, frog, and sea urchin, and among other SET domain proteins from these species. Asterisks mark where residues are identical across all twelve proteins. C) Modeling of Set8 with SAH and peptide from H4 bound to the enzyme. Shown are representative structures after 500ns of molecular dynamics for Set8, Arg634Gly and Arg634Gly, His638Leu mutations in Set8. D) Total length of time during 500 ns simulations that ligands remained in binding pocket as measured by distances between key atoms. The two distance measurements shown were selected because they were the most stable interactions between Set8 and H4 peptide and between Set8 and SAH. The selected hydrogen bond to the peptide was also the most stable interaction with the peptide and one of the last to be broken. Circles represent values from four replicate simulations. E) Western blot of third instar larval brain extracts from Oregon R wild type and the indicated Set8 mutants using anti-Set8 and anti-β-tubulin antibodies. F) Quantification of anti-Set8 signal on western blots by densitometry (see methods). Shown is the mean and standard deviation of measurements (circles) from technical replicates across four biological replicates. Oregon-R normalized signal was set to 1 for each replicate. Significance was determined by a one-way Anova followed by Tukey’s multiple comparison test. **** indicates p<.0001 and ns indicates not significant.

We inserted *Set8*^*RG*^ and *Set8*^*RGHL*^ transgenes into the same chromosomal landing site used for the KMT5A rescue experiments (Figure 2A). We then assessed expression of these transgenes in a *Set8*^*20/20*^ background by immunoblot analysis of third instar larval brain extracts. As demonstrated previously, there is no detectable Set8 protein in *Set8*^*20/20*^ homozygous null mutant animals^40^ (Figure 3E). The Set8(RGHL) mutant protein accumulates to about 10% of Oregon-R wild-type control (Figure 3E, F), suggesting that binding of the SAM cofactor stabilizes Set8 protein. Consistent with this result, *Set8*^*RGHL*^ animals are phenotypically similar to *Set8*^*20/20*^ null mutants, arresting development as early pupae (Figure 4A, B). Interestingly, *Set8*^*RGHL*^ wandering larvae accumulate melanotic masses that we did not observe in *Set8*^*null*^ animals. This phenotype is associated with immune response and was previously reported to be variably expressive and penetrant in both *Set8*^*20*^ and *Set8*^*1*^*/Df(3R)red3l* animals.^55^ In contrast, levels of the Set8(RG) missense protein are comparable to those of wild type Set8 expressed from a control *Set8*^*WT*^ transgene (Figure 3E, F), indicating that the R634G mutation does not impact protein stability. The *Set8*^*RG*^ transgene rescues the early pupal lethality observed in *Set8*^*20/20*^ mutants, but only ∼50% of the *Set8*^*RG*^ animals eclose as adults compared to the *Set8*^*WT*^ control (Figure 4A, B). A majority of *Set8*^*RG*^ mutant flies had rough eyes (86%, Figure 4C), as was previously shown for flies harboring the *Set8*^*1*^ hypomorphic mutation, which is caused by a P-element insertion in the 5’-UTR.^27,40,61^ *Set8*^*RG*^ also behaves as a hypomorphic allele, as animals containing two *Set8*^*RG*^ transgenes have a less severe phenotype than those containing one (Figure 4B). These data indicate that the *Set8*^*RG*^ allele is not null, and thus Set8(RG) protein retains at least some Set8 function *in vivo*.

**Figure 4.**
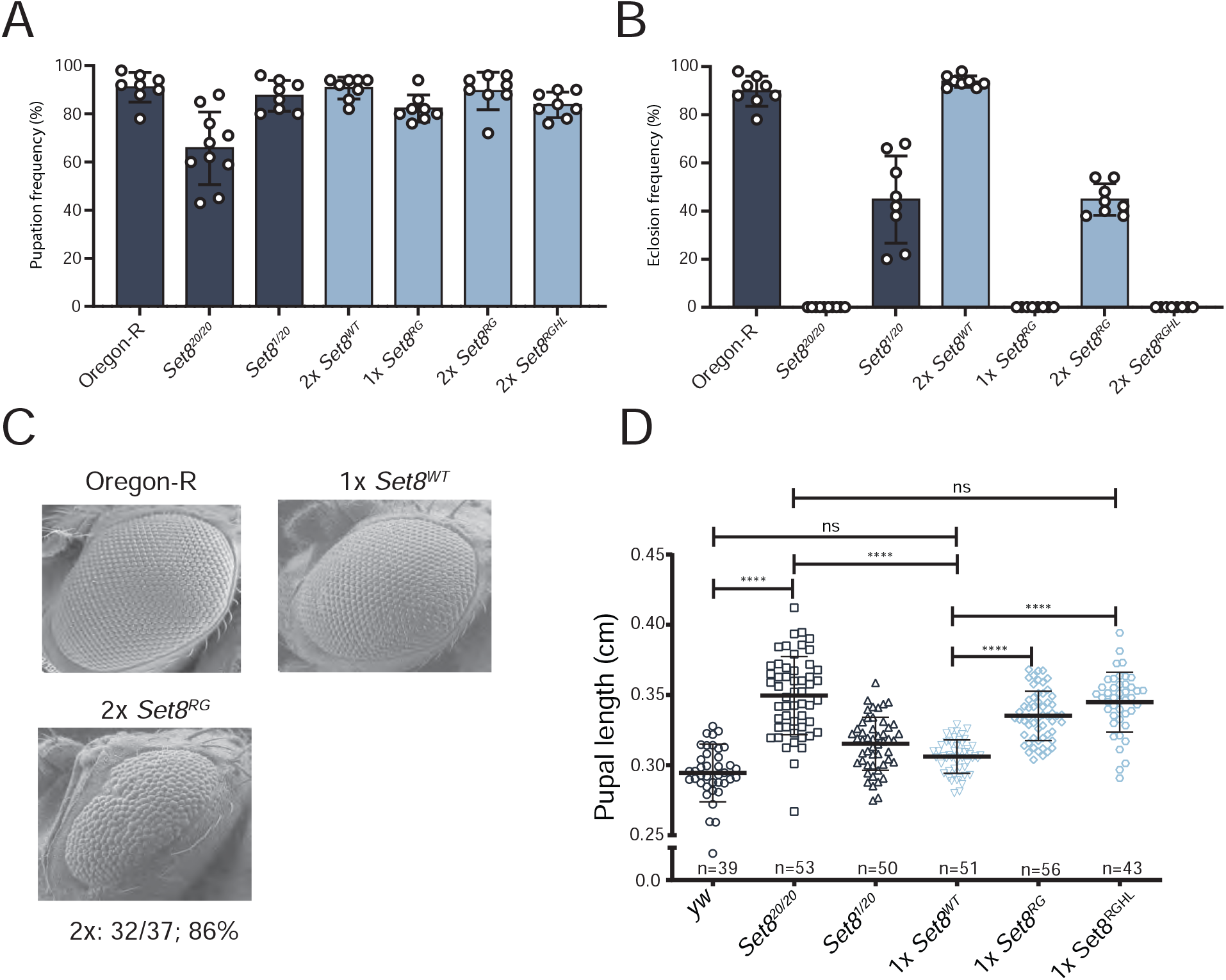
The *Set8*^*RG*^ mutant phenotype is not null. A) Pupation and B) Eclosion into adults of *Set8*^*RG*^ and *Set8*^*RGHL*^ mutants. Each circle represents the percentage of 40-50 larvae in a vial that reached pupation or adulthood. The mean and standard deviation of these percentages for 8 vials are shown for the indicated genotypes. Note that the Oregon R and *Set8*^*20/20*^ data in panels A and B are identical to Figure 2B and 2C, respectively, and shown here to allow comparison. C) SEM images of adult eyes of flies of the indicated genotypes. Penetrance and transgene copy number are as in Figure 2 legend. D) Pupal length was measured for animals of the indicated genotypes. Each symbol represents a single pupa. Thick bar indicates the mean and thin bars indicate standard deviation. ns indicates not significant and **** indicates p < 0.0001 by Student’s t test.

To further characterize the *Set8*^*RG*^ mutant, we more closely evaluated the process of pupariation in our collection of *Set8* mutants. Easily recognizable developmental events occur during the larval to pupal transition in *Drosophila*, including eversion of the anterior spiracles and gas bubble translocation from the posterior to anterior end of the pupa. Whereas S*et8*^*+/20*^ heterozygotes and *Set8*^*1/20*^ hypomorphs progress normally through these developmental milestones, *Set8*^*20/20*^ mutants fail to complete both anterior spiracle eversion and gas bubble translocation, resulting in pupae with increased length compared to control and *Set8*^*1/20*^ hypomorphs (Figure 4D). *Set8*^*RG*^ animals displayed a slight defect in completion of these pupariation events compared to *Set8*^*WT*^ control animals. Pupariation defects observed in *Set8*^*RGHL*^ animals were like those in *Set8*^*20/20*^ mutants. Interestingly, *Set8*^*RG*^ mutants also displayed a slight increase in pupal length compared to *Set8*^*WT*^ controls that did not reach the severity observed in *Set8*^*20/20*^ mutants (Figure 4D). These data demonstrate that the SET catalytic domain mutant *Set8*^*RG*^ displays intermediate pupariation defects between wild type and null alleles of Set8, suggesting that successful completion of the larval to pupal transition may require both catalytic and non-catalytic functions of Set8.

### Mutants of H4K20 and Set8 are phenotypically distinct

Mutation of lysine methyltransferases can result in disruption of multi-protein complexes, causing pleiotropic phenotypes independent of histone methylation.^20,31,45,82,87,107^ In addition, Set8 has non-histone substrates and non-catalytic functions.^23,26,34,76,80,83,100,108^ Thus, one cannot conclusively determine functional roles for H4K20me solely by mutating Set8. Another genetic strategy to address the contribution of H4K20me to various genomic processes is to change H4K20 to a residue that cannot be modified by Set8. However, this genetic strategy is not usually employed in metazoan systems because in these organisms the replication-dependent (RD) histones (H1, H2A, H2B, H3, and H4) are encoded by multiple genes located at different loci, making genetic manipulation extremely difficult. In contrast, in *Drosophila melanogaster* all ∼100 replication-dependent histone genes are tandemly arrayed at a single locus that can be removed with a single genetic deletion. The early developmental arrest caused by homozygosity of this deletion can be rescued with a single, ectopic transgene encoding 12 tandemly arrayed histone wild type gene repeats (HWT; Figure 5A, see Meers et al. 2018^52^ for details on array construction). This strategy allows us to engineer histone genotypes encoding mutant histone proteins in which a given residue is changed to one that is not a substrate for its cognate modifying enzyme.^5,41,49–52,65,66^

**Figure 5.**
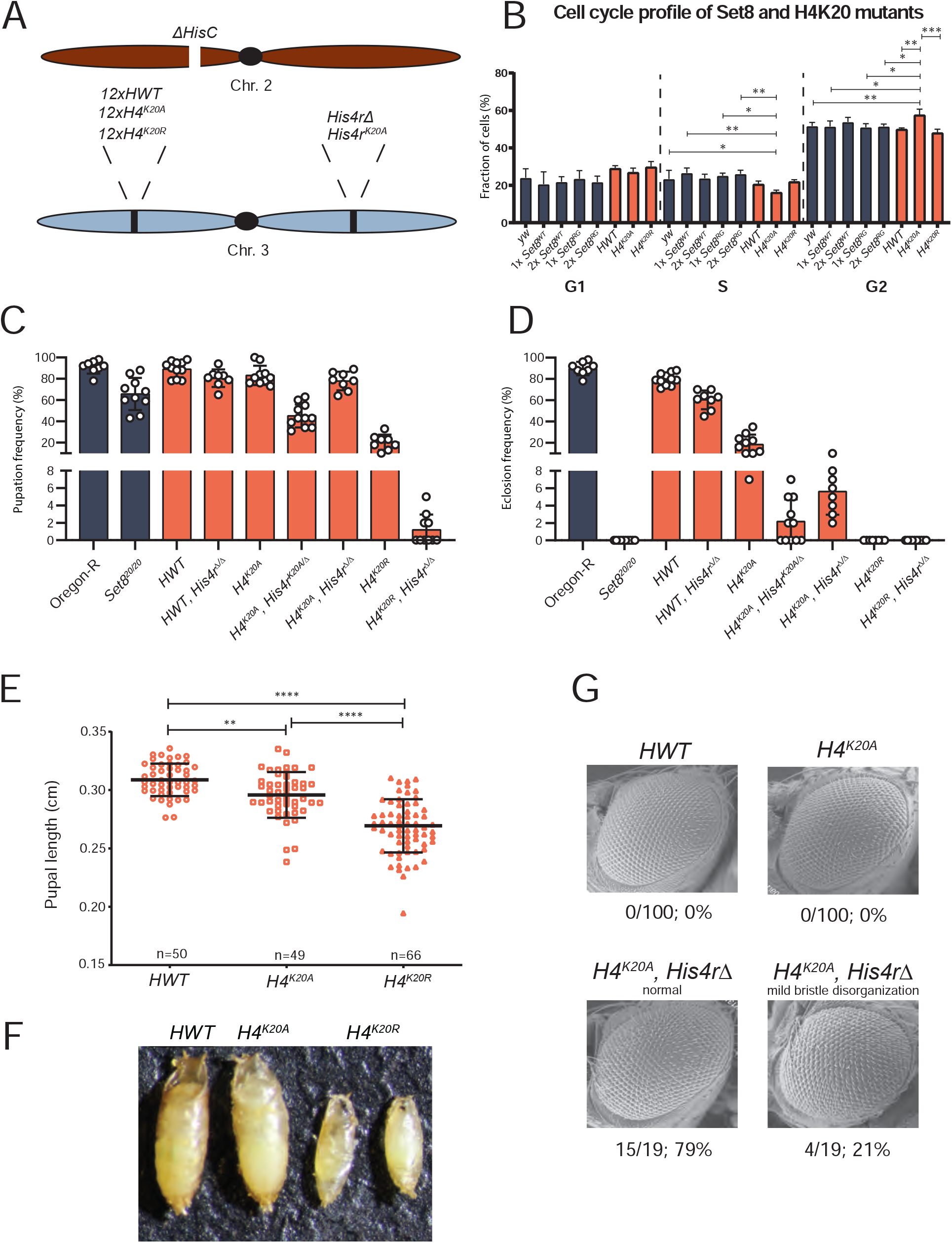
H4K20 mutant phenotypes differ from Set8 mutant phenotypes. A) Diagram of histone mutant genotypes. A deletion of HisC on the second chromosome is rescued by a third chromosome containing a transgenic 12x histone gene arrays and either with or without a mutation of *His4r*. B) FACS analysis of DNA content within cells obtained by dissociation of larval wing imaginal discs. The percentage of cells in each phase of interphase is shown for the indicated genotypes. C) Pupation and D) eclosion into adults of different H4 mutants. Each circle represents the percentage of 40-50 larvae in a vial that reached pupation or adulthood. The mean and standard deviation of these percentages for 8-11 vials are shown for the indicated genotypes. E) Pupal length was measured for animals of the indicated control and H4 genotypes. Each symbol represents a single pupa. Thick bar indicates the mean and thin bars indicate standard deviation. ** indicates p<0.004 and **** indicates p < 0.0001 by Student’s t test. F) Representative image used for the pupal length data in panel E. G) SEM images of adult eyes of flies of HWT control and the indicated H4 mutant genotypes. Rough eye phenotype penetrance is indicated below each image.

Using this strategy, we demonstrated previously that *H4*^*K20A*^ mutant animals can survive to adulthood^49^ (Figure 5C, D). By contrast, 100% of *Set8*^*20/20*^ null animals die as larvae or early pupae^40^ (Figure 2C). This stark phenotypic difference between *H4*^*K20A*^ and *Set8* mutants suggests that certain *Set8* phenotypes might not be due to loss of H4K20me, but rather to loss of methylation of its non-histone substrates or non-catalytic functions. To investigate this disparity further, we first generated *H4*^*K20R*^ mutants^52^ and compared the resulting phenotypes to those of *H4*^*K20A*^ animals as well as of *Set8* mutants. Whereas a fraction of *H4*^*K20A*^ mutants can survive to adulthood, we found that all *H4*^*K20R*^ mutants fail to eclose as adults, although some reach the pharate adult stage (Figure 5C, D). In addition, *H4*^*K20R*^ animals pupate much less frequently than either *H4*^*K20A*^ mutants or *H4*^*HWT*^ controls. Notably, the *H4*^*K20R*^ mutant pupae are much smaller and shorter than either *H4*^*HWT*^ control or *H4*^*K20A*^ mutant pupae, indicating a growth defect (Figure 5E, F). Despite this defect, we did not detect a change in cell cycle progression by FACS analysis of cells from *H4*^*K20R*^ wing imaginal discs (Figure 5B). In contrast, *H4*^*K20A*^ cells accumulate in G2 relative to controls, with a concomitant reduction in S phase (Figure 5B). Notably, Set8 deficient cells arrest in G2/M in both flies and mammalian cell culture.^14,40^ Taken together with the overall eclosion frequency differences, these data demonstrate that the *H4*^*K20R*^ mutation is more severe than the *H4*^*K20A*^ mutation developmentally, but that each mutation influences cellular mechanisms in unique ways (See Discussion).

One complication of these studies is that the fruitfly genome contains a single-copy replication-independent H4 gene (*His4r*) on chromosome 3 (i.e., located outside of the RD histone gene array on chromosome 2). *His4r* encodes an H4 protein that is identical to the RD H4^3^. Although this gene is non-essential (Figure 5C, D), we and others have found that His4r can partially compensate for loss of RD H4.^5,19,28,49^ Therefore, we used CRISPR-Cas9 to engineer two *His4r* alleles (a deletion^5^, *His4r*^*Δ* 5^ and a K20A mutant, *His4r*^*K20A*^) and we combined them with the appropriate RD histone mutant genotypes (Figure 5A). As shown in Figures 5C and D, homozygous loss of *His4r* in an *H4*^*K20A*^ background (*H4*^*K20A*^, *His4r*^*Δ/Δ*^) reduces viability, but does not eliminate it, indicating that His4r expression is important for the observed viability of *H4*^*K20A*^ mutants but is not required. Expressing one copy of *His4r*^*K20A*^ further reduces viability (Figure 5C, D), suggesting a dominant toxicity of the H4K20A protein. In contrast, deleting *His4r* in an *H4*^*K20R*^ background did not appreciably change the lethal period of *H4*^*K20R*^ animals (Figure 5C, D).

We next compared H4K20 and Set8 mutant phenotypes, focusing on pupariation and eye development. In contrast to *Set8*^*20/20*^ null mutants, which display defects during pupariation, >80% of the *H4*^*K20A*^ and *H4*^*K20R*^ animals complete proper anterior spiracle eversion and gas bubble translocation. Similarly, the viable *Set8*^*RG*^ and *Set8*^*1/20*^ mutants did not exhibit defects in anterior spiracle eversion or gas bubble translocation. Both *Set8*^*RG*^ and *Set8*^*1/20*^ mutants have rough eyes^27^ (Figure 4C), indicating that Set8 is required for eye development. In contrast, none of the *H4*^*K20A*^ mutants had rough eyes when *His4r* was present, whereas ∼21% of *H4*^*K20A*^, *His4r*^*Δ/Δ*^ animals only had mild disorganization of interommatidial bristles (Figure 5G). These results suggest that the roles of Set8 and H4K20me in eye development are distinct, and further highlight that the differential effects of Ala and Arg substitutions at H4K20. We conclude that H4K20me does not mediate all functions of Set8 because mutating H4K20 and Set8 cause different developmental phenotypes.

## Discussion

We use genetic and genomic approaches in *Drosophila* to investigate how histone PTMs, and the enzymes that install them, contribute to animal development. It is particularly informative to determine where these contributions differ. Our results indicate that only a subset of the essential functions of the H4K20 mono-methyltransferase, Set8, are mediated by H4K20me. The data also reveal that, although H4K20me is formally dispensable for completion of development, the lysine residue nonetheless plays an important role.

### *Drosophila* Set8 and human KMT5A are orthologous

We showed that human KMT5A can substitute for all Set8 functions during *Drosophila* development, except in the eye, where we observe a minor disruption in ommatidial organization that manifests as a rough eye in *KMT5A*-rescued adults. The eye phenotype likely does not result from changes in methylation of substrates, as we found that the human KMT5A SET domain can fully substitute for that of Set8, even in the eye. Rather, full developmental eye function is instead provided by the non-catalytic amino terminal 339 amino acids of Set8, which is conserved in other Diptera, but not in humans or other vertebrates and invertebrates. Nonetheless, we found that the rough eye phenotype was more penetrant in *Set8*^*ΔN*^-rescued animals than it was in the *KMT5A*-rescued animals. We designed Set8^ΔN^ to be the same size as KMT5A and retain the conserved PIP degron and Cdk consensus phosphorylation site^96^ found in KMT5A, which therefore might provide some function during eye development. Because KMT5A can perform nearly all the biological functions of Set8 in *Drosophila*, studies of Set8 could be applicable to human biology and disease, particularly because aberrant levels of KMT5A are implicated in the development of and increased risk in certain breast, brain, and liver cancers.^18,22,54,59,76,83,97,104,106^ KMT5A has also been shown to regulate androgen receptor-mediated transcription in prostate cancer.^99^

### Mutation of the SET domain does not abolish *in vivo* function of Set8

Whether methyltransferase activity is required for all the cellular and developmental roles of SET domain proteins remains an open question in the field. This question is generally addressed by testing the *in vivo* function of “catalytically dead” enzymes. Previous *in vitro* studies showed that an R265G mutation eliminates catalytic activity of KMT5A.^60^ We found that the corresponding *Set8*^*R634G*^ mutation does not cause a null mutant phenotype and supports development into viable adults. This result suggests that methylation of both H4K20 and non-histone substrates of Set8 is not required for completion of development in *Drosophila*. However, we do not know whether *Set8*^*RG*^ flies retain some H4K20 methylation. Thus, one possibility is that an enzyme other than Set8 could methylate H4K20 in *Set8* mutant animals, but the levels of H4K20me attained would not provide full biological function. Another possibility is that the *Set8*^*RG*^ mutant is a catalytic hypomorph *in vivo*. Consistent with this possibility, the *Set8*^*RG*^ mutant phenotype resembles that of the previously described *Set8*^*1*^ hypomorphic mutant (viable with rough eyes),^27,40^ and we found that two copies of the *Set8*^*RG*^ transgene provide more function than one copy, indicating that *Set8*^*RG*^ is a genetic hypomorph. In addition, our structural analyses revealed that R634G disrupts interactions within the SAM binding domain but does not eliminate the possibility that SAM and K20 might still occupy the active site of the enzyme, albeit less avidly. Notably, other studies have concluded that catalytic activity is not required for *in vivo* function of the H3K4 mono-methyltransferase (Mll3/4, Trr).^25,68^ Thus, critical non-catalytic roles of histone methyltransferases may be the norm rather than the exception.

### Comparative genetic analyses support distinct developmental roles for Set8 and H4K20me

Our analysis of H4K20 mutants is consistent with the idea that Set8 provides essential functions during metazoan development that do not include H4K20 methylation. Animals entirely lacking H4K20me (*H4*^*K20A*^, *His4r* ^*Δ /Δ*^ and *H4*^*K20A*^, *His4r*^*K20A/Δ*^) can develop into adults with no obvious morphological defects, whereas all animals lacking the H4K20 methyltransferase (*Set8*^*20/20*^) die in early pupal stages. This difference in phenotype supports the hypothesis that *Set8*^*20/20*^ inviability is due, at least in part, to loss of non-histone substrate methylation and/or non-catalytic functions of Set8. Nevertheless, H4K20 is clearly quite important, as only a small fraction of *H4*^*K20A*^ mutants complete development and *H4*^*K20R*^ mutants are inviable. Moreover, ectopic expression of H4K20A mutant histones in cultured human cells supports a role for H4K20me in S phase progression, particularly in late replicating heterochromatin.^13^

The phenotypic differences we observe between *H4*^*K20A*^ and *H4*^*K20R*^ mutants are intriguing, as both substitutions are expected to eliminate H4K20me. The differences may well be attributable to idiosyncratic structural properties of H4A20-vs. H4R20-containing nucleosomes, relative to wild type. In particular, the side chains of Alanine and Arginine differ in both size and charge and thus may differentially impact interaction of the H4 tail with chromatin binding complexes irrespective of H4K20 methylation. For instance, proteins that bind unmethylated H4K20 (BRCA1-BARD1) do not recognize H4K20A nucleosomes.^58^ Given the proximity of H4K20 to the nucleosome core, these mutations may variably influence chromatin structure, or affect the modification of other residues on the H4 tail or on other histones within the nucleosome. Notably, the assumption that a Lys for Arg substitution would be less detrimental than a Lys for Ala substitution (because Lys and Arg have a similar side chain structure and are both positively charged) is not born out by our data. Regardless of the precise mechanism, our genetic analyses provide important insight into H4K20me function *in vivo*, and suggest that future biochemical, proteomic, and ultrastructural studies of these histone mutants will be informative.

## Materials and Methods

### Fly stocks and husbandry

Fly stocks were maintained on standard corn medium with molasses provided by Archon Scientific (Durham). The *Set8*^*20*^ stock used in this study was a generous gift from Ruth Steward. The *Set8*^*1*^ (#10278) stock was obtained from the Bloomington Stock Center.

### Set8, KMT5A, and chimeric transgenes

For the *Set8*^*WT*^ transgene a 5493 bp genomic fragment was amplified from a wild type fly extract using the following primers 5’ acttatacacttcattcct 3’ and 5’ tacccgcctgatgcgaattt 3’. The genomic fragment was cloned into pDEST w+ attB (Supplemental Figure 2). *Set8*^*RG*^ and *Set8*^*RGHL*^ were constructed using site-directed mutagenesis using the Q5 Site-directed Mutagenesis kit on pDEST w+ attB Set8^WT^ (NEB E0554S). KMT5A, Set8^ΔN^, N-KMT5A::Set8-C, and N-Set8::KMT5A-C sequences were synthesized using GENEWIZ gene synthesis (Supplemental Figure 3) and cloned into pDEST w+ attB digested with AgeI and MluI (Supplemental Figure 2). Transgenes were sequence-verified and injected into VK33 on chromosome 3L and screened for positive transformants by BestGene. Recombinant flies were generated by crossing transgenic flies with flies containing *Set8*^*20*^ and screening single F2 male progeny for the presence of both the appropriate transgene and *Set8*^*20*^.

### Western blots

Twenty brains from third instar wandering larvae of each genotype were collected in 1xPBS (137 mM NaCl, 2.7 mM KCl, 10 mM Na_2_HPO_4_, 1.8 mM KH_2_PO_4_). 1xPBS was removed, 100 uL of RIPA buffer (50 mM Tris pH 7.5, 0.1% SDS, 0.5% Sodium Deoxycholate, 1% NP-40, 150 mM NaCl, 5 mM EDTA)) was added to each sample. Larvae were homogenized in RIPA buffer using a pestle and incubated on ice for 30 minutes. Samples were then centrifuged for 15 minutes at top speed at 4C. Supernatant was separated from pellet and protein concentration was assessed using a Bradford assay. Then 4x Laemmli sample buffer (BioRad 1610747) with 10% β-mercaptoethanol was added to each sample at a 3:1 ratio. Samples were boiled for 10 minutes and equal protein (∼10 ug) was loaded on a 12% SDS-PAGE gel. Proteins were transferred to a 0.45nm nitrocellulose membrane for 60 minutes at 100V. Membranes were blocked with 5% milk in 1xTBS-Tween (10mM Tris, 150 mM NaCL, 0.1% Tween20) for 60 minutes then blotted with primary antibodies (Set8: Novus Biologicals 44710002; β-tubulin: Abcam ab6046) in 5% milk in 1xTBS-Tween overnight. Blots were quickly washed 3x then for 10 minutes 3x. Blots were incubated with secondary antibody (goat anti-Rabbit) in 5% milk in 1xTBS-Tween for two hours at room temperature. Blots were again quickly washed 3x then for 10 minutes 3x. Blots were then incubated with SuperSignal™ West Pico PLUS Chemiluminescent Substrate (Thermo Scientific 34580) and imaged using a GE Amersham Imager. Quantification was performed using FIJI. Briefly, the signal of each band on the Set8 blot and β-tubulin blot was quantified using a box of equal area. Signal from *Set*^*20/20*^ was subtracted from each Set8 value, then divided by the corresponding β-tubulin signal for each lane. Finally, the value of *Oregon-R* was set to 1, so values of all other genotypes are relative to that genotype.

### Viability assays

To investigate the requirement of Set8 and H4K20me for organismal viability, we enriched cultures of each genotype for 1^st^ instar larvae by manually separating them from their wild type siblings and monitored survival to pupal and adult developmental stages. Mean pupation and adult values and pairwise comparisons for each genotype can be found in Supplemental Figure 4. Crosses to generate histone mutant genotypes were the same as previously reported.^4,52^

### CRISPR for *His4r*

The *His4rΔ* allele utilized in this study was the same generated by Armstrong et al. 2018. Here we generated a point mutation allele (*His4r*^*K20A*^) using CRISPR-Cas9 mutagenesis. The genomic region including *His4r* was amplified using the following primers 5’-gctgcgccgttagataaagc-3’ and 5’-agcaatcggagtccatg-3’ and TOPO cloned in pENTR. The codon for His4r^K20^ was changed to Ala using the Q5 Site-directed Mutagenesis kit (NEB E0554S). The same gRNA constructs in pCFD3 from Armstrong et al. 2018 were co-injected with the K20A-mutated *His4r* repair construct into *Drosophila* embryos expressing Cas9 from the nanos promoter. Positive hits were screened using a BbsI site created by the Lys to Ala mutagenesis.

### Scanning electron microscopy

Flies were deyhydrated in ethanol and images of compound eyes were taken using a Hitachi TM4000Plus table top SEM microscope at 10kV and 150x magnification.

### FACS

Wing imaginal disc nuclei from third instar wandering larvae of each genotype were sorted into G1, S, and G2 populations by a FACSAria II or III based on DAPI intensity as previously described.^5,53^

### Protein sequence analyses

Figure 1A: PRDM and SET domain methyltransferase protein sequences (Supplemental Figure 5) were compiled and aligned with ClustalOmega using the msa package.^11^ A distance matrix was calculated by identity using dist.alignment in the seqinr package. A phylogenetic tree based on the distance matrix was generated and then plotted using ggtree.^101–103^

Figure 2C: The full-length *Drosophila melanogaster* Set8 protein sequence was BLASTed against the refseq_protein database using the default parameters. The top 1000 hits were compiled and manually sorted to include only one protein isoform per organism. Proteins with percent identities to the full-length *Drosophila melanogaster* Set8 less than 50% were discarded. Human and mouse KMT5A proteins were retained for downstream analysis despite having percent identities lower than 50%. The remaining protein sequences (Supplemental Figure 6) were aligned with ClustalOmega from the msa package.^11^ Phylogenetic classification of each protein was performed with the taxize package and merged with the alignment information. Proteins with incomplete classification information were discarded. A phylogenetic tree was generated using the classification information and plotted using ggtree.^101–103^ The alignment of all remaining proteins was plotted in order of the phylogenetic tree and each position in the alignment was colored based on whether it matched the residue in the corresponding position of Drosophila melanogaster Set8 (blue), Human KMT5A (pink), both *Drosophila melanogaster* Set8 and KMT5A (maroon), or neither *Drosophila melanogaster* Set8 and KMT5A (black). Gaps in the alignment are represented by white space.

### Molecular dynamics simulations

Structural models of *Drosophila* WT and mutant Set8/KMT5A in ternary complexes with SAH (S-adenosyl-L-homocysteine) and H4 peptide were built using the crystallographic structure of human KMT5A in ternary complex (PDB ID: 1zkk) [PMID 15933070] as template. These structural models were then used as starting structures for molecular dynamics simulations. Four replicate explicit solvent simulations with the same starting conformations but different velocity distributions were completed for WT and each mutant using the Amber v18 software package.^21^ LEaP from the Amber software package was used to generate the explicit solvent systems in an octahedral box with charge neutralization while the GPU version of PMEMD was used to complete the simulations.^30,73^ The ff14SB force field was used for parameterization.^47^ A total of 5,000 steps of minimization were completed, followed by 500 psec heating with an NVT ensemble, and then density equilibration over 500 psec with an NPT ensemble. The production run was in the NPT ensemble for a total of 500 nsec. During the production run, Langevin dynamics with a collision frequency of 1.0 psec^-1^ was used for temperature regulation. A Berendsen barostat with a relaxation time of 1.0 psec was used for pressure regulation. The time-step was 2 fsec with hydrogen atoms constrained by SHAKE. Trajectories were analyzed for the distance between atoms in Set8/KMT5A and atoms in either the H4 peptide or SAH.

### Full genotypes of strains used in this study

*Set8*^*WT*^: *y-w-;;{Set8*^*WT*^*}, Set8*^*20/20*^

*Set8*^*RG*^: *y-w-;;{Set8*^*RG*^*}, Set8*^*20/20*^

*Set8*^*RGHL*^: *y-w-;;{Set8*^*RGHL*^*}, Set8*^*20/20*^

*KMT5A: y-w-;;{KMT5A}, Set8*^*20/20*^

*Set8*^*ΔN*^: *y-w-;;{Set8* ^*Δ1-339*^*}, Set8*^*20/20*^

*N-KMT5A::Set8-C: y-w-;;{KMT5A*^*1-214*^ *– Set8*^*555-691*^*}, Set8*^*20/20*^

*N-Set8::KMT5A-C: y-w-;;{Set8*^*1-554*^ *– KMT5A*^*215-352*^*}, Set8*^*20/20*^

*HWT: y-w-; ΔHisC; {12xHWT}*

*HWT, His4r*^*Δ/Δ*^: *y-w-; ΔHisC; {12xHWT}, His4r*^*Δ/Δ*^

*H4*^*K20A*^: *y-w-; ΔHisC; {12xH4*^*K20A*^*}*

*H4*^*K20R*^: *y-w-; ΔHisC; {12xH4*^*K20R*^*}*

*H4*^*K20A*^, *His4r*^*Δ/Δ*^: *y-w-; ΔHisC; {12xH4*^*K20A*^*}, His4r* ^*Δ / Δ*^

*H4*^*K20A*^, *His4r*^*K20A/Δ*^: *y-w-; ΔHisC; {12xH4*^*K20A*^*}, His4r*^*K20A/Δ*^

*H4*^*K20R*^, *His4r*^*Δ/Δ*^: *y-w-; ΔHisC; {12xH4*^*K20R*^*}, His4r* ^*Δ/Δ*^

## Data availability

Strains and plasmids available upon request. The authors affirm that all data necessary for confirming the conclusions of the article are present within the article, figures, and tables.

## Acknowledgements

A.T.C. and R.L.A. were supported in part by NIH predoctoral traineeships, T32-GM007092. S.K. was supported in part by an NIH diversity supplement to grant R01-DA036897 (to R.J.D., A.G.M. and B.D.S.) and by a postdoctoral traineeship, T32-CA009156. This work was supported by NIH grants R35-GM136435 (to A.G.M) and R01-GM124201 (to R.J.D.). We thank Ruth Steward for generously providing fly stocks and Megan Butler for critical reading of the manuscript.

